# Investigating Cement-Based Surfaces as a Sustainable Flooring Solution to Improve *Ascaris* Egg Removal and Inactivation in Low-Resource Settings

**DOI:** 10.1101/2025.02.18.638796

**Authors:** Claire E. Anderson, Suhi Hanif, Jason Hernandez, Yoshika Crider, Michael Lepech, Sarah L. Billington, Alexandria B. Boehm, Jade Benjamin-Chung

**Affiliations:** Department of Civil and Environmental Engineering, Stanford University, Stanford, CA, United States of America; Department of Epidemiology and Population Health, Stanford University, Stanford, CA, United States of America; King Center on Global Development, Stanford University, Stanford, CA, United States of America; Chan Zuckerberg Biohub, San Francisco, CA, United States of America

## Abstract

Soil-transmitted helminths, like *Ascaris*, are significant contributors to disease burden in low- and middle-income countries (LMICs). Infections are associated with growth faltering and mortality in children and are often transmitted through contact with eggs in fecally contaminated soil. Interventions, like replacing household soil floors with cement-based alternatives, may reduce exposure to *Ascaris* eggs, but there are currently no estimates on the removal or survival of *Ascaris* eggs on cement-based surfaces. This study addresses that knowledge gap by evaluating the removal of *Ascaris* eggs from mopping and the survival of *Ascaris* eggs on two cement-based mixes: an Ordinary Portland Cement (OPC) mortar and an OPC mortar with fly ash, which provides a more sustainable alternative to the OPC mortar mix. We assessed egg survival at two temperatures representing the dry (15°C) and wet (34°C) seasons in Bangladesh using two different egg enumeration methods. After mopping, over 92% of viable eggs were removed from surfaces, with no significant differences between cement-based mixes (p = 0.51). The first-order decay rate constants (*k*) of *Ascaris* eggs were similar between mix designs (p = 0.62) but varied significantly between temperatures (p = 4.2 x 10^-25^) and egg enumeration methods (p = 2.4 x 10^-8^). The *k* values were of greater magnitude at 34°C compared to at 15°C. At 15°C, *k* values were not significantly different from zero, indicating no inactivation. The *k* values we obtained were comparable to those reported in previous studies for different matrices, indicating comparable inactivation of *Ascaris* eggs on cement-based surfaces compared to liquid and semi-solid matrices. These results provide some of the first estimates of removal efficiencies and inactivation times in realistic environmental conditions for *Ascaris* on surfaces while supporting the use of OPC mortar mix designs with fly ash in interventions to reduce *Ascaris* transmission in rural LMIC households.

**Author Summary:** Soil-transmitted helminths, like *Ascaris*, are parasites that are major contributors to disease in children and women of childbearing age in low- and middle-income countries. Interventions, like replacing soil floors in households with cement-based flooring, may reduce exposure to *Ascaris* eggs which cause infections, but there is little information on how or why these interventions may be effective. This study investigates the effectiveness of simple cleaning methods, like mopping, in removing *Ascaris* eggs from cement-based surfaces and explores how long these eggs can survive on these surfaces under different environmental conditions. We tested two types of cement-based surfaces, a traditional cement-based mortar mix, and a more sustainable cement-based mortar mix, and found that mopping removed over 92% of Ascaris eggs, with no differences between cement mixes. Experiments simulating wet and dry seasonal conditions showed that Ascaris eggs survive longer in cooler environments, again with no differences between cement mixes. These findings provide important insights into the role of cement-based flooring in interrupting disease transmission and suggest sustainable cement-based mortar mixes are a feasible alternative to traditional cement-based mortar mixes.

## Introduction

Soil-transmitted helminths infect approximately 24% of the global population, but infections are most common in low- and middle-income countries (LMICs) [1]. Prevalent soil-transmitted helminths include *Ascaris lumbricoides* (roundworm), *Trichuris trichiura* (whipworm), and *Necator americanus* (hookworm) [1,2]. *Ascaris* is a particular concern in Bangladesh, as it is endemic to all 64 districts, and poses the largest risk to preschool-age children, school-age children, and women of childbearing age [3]. In 2010 in Bangladesh, 79.8% of school-age children were infected with one or more helminth species [4]. For *Ascaris*, the most intense infections occur in school-age children and can cause malnutrition, growth stunting, and cognitive deficits [3,5].

Soil floors are common in LMICs [6] and are reservoirs of many pathogens, including soil-transmitted helminths. In studies that collected soil from households to test for soil-transmitted helminths, *Ascaris* eggs were consistently the predominant parasite detected [7,8]. *Ascaris* eggs contaminate soil through the feces of infected individuals, and once eggs are present in soil they can survive for up to 10 years, withstanding extreme weather conditions, and are virtually impossible to remove from soil [2,9]. After fertilized eggs embryonate in the soil, if they are consumed they can infect an individual and continue the pathogen’s life cycle. Children in LMICs are at high risk of exposure to soil-borne pathogens, including soil-transmitted helminths, as they can ingest soil during hand-to-mouth contact [10–15].

Deworming treatments are effective in the short term, but rapid reinfection occurs when environmental reservoirs, such as contaminated soil, are not eliminated. Within 6 months it is estimated that 68% of those treated become reinfected with *Ascaris* [16]. Ongoing mass school-based drug administration has occurred in Bangladesh since 2008 to reduce *Ascaris* infections, but prevalence is still over 20% in 24 of 64 districts [3,17]. In principle, water, sanitation, and hygiene (WASH) interventions could reduce contamination of soil with *Ascaris* eggs; however, studies investigating the impact of WASH interventions have had inconsistent effects on pathogen removal and disease reduction [8,18–23]. This may be due to a lack of coverage of WASH interventions, inconsistent use of interventions, or because interventions do not sufficiently prevent eggs from reaching soil reservoirs [20,22].

Replacing soil floors with finished flooring, like cement-based floors, may reduce the exposure to soil-transmitted helminths in the home environment by removing a key environmental reservoir. In contrast with WASH interventions, which may require a high level of compliance from communities, intervention uptake is nearly guaranteed with cement-based floors. Observational studies have noted that finished flooring is associated with a lower prevalence of soil-transmitted helminths [24–27], but the mechanisms of disease reduction remain unclear. An ongoing randomized trial in rural Bangladesh, Cement-based flooRs AnD chiLd hEalth (CRADLE, NCT05372068), is currently assessing whether transitioning from soil to cement floors can reduce soil-transmitted helminth infections and diarrhea in children under two years of age [28].

If cement-based floors were to be constructed as a health intervention at scale, they could contribute to significant CO_2_ emissions, as cement production contributes an estimated 5 to 10% of total global anthropogenic CO_2_ emissions [29,30]. Using an alternative cement mix, such as a mix that replaces a portion of cement with fly ash, would offset CO_2_ emissions for the subset of cement replaced [31–38]; fly ash is a by-product of coal combustion in power plants and is readily available in LMICs. Although fly ash may contain heavy metals [39], these metals become largely immobilized in the cement matrix after curing through absorption and chemical reactions [40,41], eliminating significant risks of exposure for household members in contact with fly ash-containing cement-based floors. While replacing a portion of cement with fly ash would offset CO_2_ emissions, it is unknown whether *Ascaris* egg survival and removal would be similar on these surfaces compared to traditional cement-based surfaces. In principle, differences in the cement-based mix’s surface roughness properties, moisture retention, and other factors, could impact the survival and removal of pathogens on each mix’s surface, as pathogens could be shielded from or made more vulnerable to environmental stressors like desiccation. Though similar structural properties have been found between traditional mortar mixes and fly ash mortar mixes [33–38], the impact of fly ash as a cement replacement in mortar mixes on the survival and removal of soil-transmitted helminths is unknown.

No prior studies have investigated the survival and removal of *Ascaris* on surfaces, nor the factors that affect their inactivation. Ammonia and 254-nm ultraviolet light can effectively remove *Ascaris* from liquid matrices [42–44], but it is unknown whether they can remove *Ascaris* from surfaces. Further, in LMICs, these disinfection products are not attainable for most rural households; instead, common floor cleaning methods include sweeping or mopping with water.

In this study, we evaluated the removal and survival of *Ascaris* eggs on cement-based surfaces while mimicking conditions in a setting where soil-transmitted helminths are common. We hypothesized that removal and survival would be similar on a typical cement mix and an alternative mix replacing some cement with fly ash. We also hypothesized that *Ascaris* first-order decay rate constants would be greater in the wet season temperature, which is warmer than the dry season temperature. The results of this study will indicate if sustainable alternatives to cement-based flooring are appropriate for large-scale infrastructure interventions to prevent the persistence of *Ascaris* in households.

## Methods

### Cement-Based Tiles

We used a traditional Ordinary Portland Cement (OPC) mortar mix to create cement-based tiles and finished the tiles with a smooth finish of cement paste. We also used an OPC mortar mix with 25% Class F fly ash as a cement replacement to create cement-based tiles and finished those tiles with a smooth cement paste finish containing the same 25% Class F fly ash replacement. In this manuscript, we will refer to the first tile type as the OPC mortar mix tiles and to the second as the OPC fly ash mortar mix tiles. Mix design details are included in the Supplemental Information (SI, Table S1).

Cement-based tiles were made according to ASTM C192 Standard Practice for Making and Curing Concrete Test Specimens in the Laboratory [45]. The two cement-based mixes used in this study were prepared in the same manner as those used in Bangladesh in the CRADLE study. The mixes were poured into 127 mm square molds with 12.7 mm depth and allowed to set. After demolding, the tiles were 7 day wet-cured in a lime bath followed by a minimum 28-day air-cure. For testing, each 127 mm square tile was delineated into four, approximately 3 cm x 3 cm, square quadrants using a wax pencil. Some tiles were used twice in experiments; before being used a second time they were disinfected by autoclaving at 121°C and washing thoroughly with water, after which they were allowed to dry at room temperature (approximately 21°C) for a minimum of 5 days before reuse in experiments.

### Ascaris suum Stock

*Ascaris suum* has been extensively used as a surrogate for *Ascaris lumbricoides* [42,46–50], as the eggs are morphologically and physiologically indistinguishable and their genomes exhibit 98.1% similarity [2,47]. *Ascaris suum* stock was purchased from Excelsior Sentinel, Inc. (Trumansburg, NY, USA). The stock solution was made up of roughly 10^6^ fertilized *Ascaris suum* eggs, derived from sieved pig feces, and 50 mL of 0.1 N sulfuric acid to prevent mold growth. Upon arrival, the egg stock solution was stored at 4°C to prevent eggs from embryonating; eggs were stored between two and seven months before use in experiments. The stock solution was sampled throughout the storage time period for egg enumeration and development assessment. Each time prior to using the stock, the solution was inverted 10 times to ensure a well-mixed solution.

The dry weight of the total solids in the stock solution was determined by spiking 25 μL of the stock solution onto a weigh boat and allowing the sample to dry for approximately 3 hours, until the stock was no longer visibly wet. The average recorded weight after drying for five samples was used to determine the dry weight of the total solids per volume of the stock.

### Outcomes

The study outcomes included 1) the percent removal of viable eggs on the two different cement-based mixes from mopping and 2) the first-order decay rate constants of the eggs (both viable eggs and the broader category of developed eggs) over time on the two cement-based mixes in high- and low-temperature conditions.

### Experimental Procedure

#### Inoculation

To inoculate the cement tiles with *Ascaris*, 25 μL of the stock solution (*Ascaris suum* eggs from sieved pig feces in 0.1 N sulfuric acid) was added to 2 mL of autoclaved DI water. The solution was spread evenly across one 9 cm^2^ quadrant of the tile using a sterile disposable plastic needle (Fisherbrand, Waltham, MA, USA) and allowed to dry for approximately 1 hour until the surface was no longer visibly wet.

#### Recovery

Eggs were recovered from the cement tile using a new method, referred to as the “washing method” in this study. With the washing method, roughly 3 mL of autoclaved DI water was added to one 9 cm^2^ quadrant of the tile, and the surface of the quadrant was agitated with a sterile disposable plastic needle (Fisherbrand, Waltham, MA, USA). The water was pipetted off the surface and added to a sterile, 50 mL centrifuge tube (CORNING, Corning, NY, USA). The process of water addition, agitation, and recovery was repeated until 15 mL of water was collected from the tile quadrant; this typically took 10 - 15 minutes to achieve per quadrant of the tile. Then, 15 mL of 0.1 N sulfuric acid (Sigma-Aldrich, St. Louis, MO, USA) was added to the tube to prevent mold growth, resulting in a total volume of 30 mL per sample. To embryonate the eggs, the sample was incubated in darkness for 32 - 35 days at 26°C. This combination of incubation time and temperature is sufficient to embryonate the maximum number of eggs in the sample [46,51,52].

#### Enumeration

Samples were centrifuged at 1000 x g for 3 minutes and 29 mL of the supernatant was removed and discarded according to biosafety guidelines. The remaining 1 mL sample was transferred to a Sedgewick-Rafter slide (Electron Microscopy Sciences, Hatfield, PA, USA) for enumeration under a microscope [53]. A microscope (Swift Optical Instruments, Inc. San Antonio, TX, USA) was used to view the slide at 10X magnification in the program ToupLight (ToupTek, Hangzhou, Zhejiang, P.R. China) with a 0.5X magnification camera attachment (OMAX Microscopes, Irvine, California, USA).

Eggs were counted and classified into one of 16 development stages [52] or as dead/non-viable. The total number of eggs includes all development stages, dead, and non-viable eggs. We determined viability through both conventional and developmental methods [52]. For the conventional enumeration method, eggs at stage 15 (which had the most well-developed larvae prior to excystation) of the developmental process were considered viable and all other eggs were considered non-viable. For the developmental enumeration method, we grouped into five categories based on their development stages: (1) Unembryonated, stage 1; (2) Embryonated, stage 2 to 7; (3) Well-developed, stage 8 to 15; (4) Excystation, stage 16; and (5) Dead or non-viable. Example photos of development categories are shown in the Supplemental Information (SI, Figure S1).

Viable *Ascaris* eggs are conventionally defined as embryonated eggs containing mobile, distinguishable larvae. Fertilized eggs become viable if they are incubated under the right conditions, while dead or unfertilized eggs cannot become viable. In conventional microscopy methods, all eggs without larvae are considered non-viable [46,52,54], however, this may undercount the total number of potentially viable eggs. For example, eggs that have larvae in well-developed stages may still be capable of later development to become fully viable. Thus, previous studies have reasoned that eggs containing mobile, distinguishable larvae as well as eggs with well-developed larvae should be considered when determining viability [46,52,54]. In removal experiments in this study, we quantified viable eggs as eggs at developmental stage 15 (conventional method). For survival experiments, we quantified viable eggs in two ways, (1) as eggs at developmental stage 15 (conventional method) and (2) as eggs in the well-developed, third category of stages (developmental method).

#### Removal Experiments

These experiments mimic *Ascaris* egg presence in the field before and after mopping on two different types of cement-based floors. On a single cement-based mix tile, each of the four quadrants on the tile was inoculated using the procedure previously described in this study. After inoculation, two of the quadrants were designated as no-mopping controls and the *Ascaris* was recovered and enumerated. The remaining two quadrants were mopped with two damp 16 cm^2^ 100% cotton cloths (Nabob Wipers, Brooklyn, NY, USA), wet with approximately 5 mL of autoclaved DI water. The two quadrants were wiped twice with side-to-side motions until the entire square was visibly damp. The remaining *Ascaris* were then recovered from the tile using the washing method. Each no-mopping control quadrant and mopping quadrant pair on the tile represents one trial. Twenty-six (26) trials were performed on tiles of each of the two cement mixes (26 trials x 2 mix design groups = 52 trials total). This n was chosen for the removal and survival experiments to provide 80% power for the main outcomes of the study. The experiments were powered to measure large effect sizes according to a unitless, standardized measure of effect size for each statistical test used [55,56]. To determine differences in the percent removal of eggs between tiles of the two cement mixes, a minimum sample size of 26 per group was required for an unpaired, two-tailed t-test.

#### Survival Experiments

These experiments mimic *Ascaris* egg survival in the field in the wet season of Bangladesh (34°C, 75% relative humidity (RH)) and dry season (15°C, 75% RH). The wet season typically ranges from April to October in Bangladesh. In Tangail, a city near the CRADLE study site, the highest normal mean monthly maximum temperature was in April, at 33.9°C, and the relative humidity was 74% [57,58]. The dry season typically ranges from November to February, and in Tanagail the lowest mean monthly minimum temperature (in January) was 11.4°C and the relative humidity was 80% [57,58]. Relative humidity was kept constant in this study through saturated salt solutions [59]. A single cement tile represented one trial and was split into four quadrants for each of the 4 time points studied (0, 2, 14, and 28 days). All four quadrants were inoculated using the procedure previously described. After inoculation, samples were sacrificially sampled at each time point, meaning that the sample was removed from the tile and was not placed back for future time points. Instead, another quadrant was sampled for later time points. Thirteen (13) trials were conducted for each cement mix at two season temperatures, the wet season and dry season (13 trials x 2 mix designs x 2 temperatures = 52 trials total). To determine differences in the first-order decay rate constants of the eggs between the two cement mixes and two temperature conditions with 80% power, a minimum sample size of 13 per group was required for an Analysis of Variance (ANOVA) with fixed effects.

#### Controls

Positive *Ascaris* egg application controls were created by spiking 0.25 μL *Ascaris* stock directly into a 50 mL centrifuge tube with 15 mL water. Then, 15 mL of 0.1N sulfuric acid was added, and the sample was incubated in darkness for 32 - 35 days at 26°C and the number of total and viable eggs was enumerated. Positive *Ascaris* application controls provide insight into the total number of eggs and the number of viable eggs applied to the tile. Positive *Ascaris* recovery controls from the cement-based tiles were created by inoculating tiles as previously described and then recovering the sample and incubating in darkness for 32 - 35 days at 26°C and enumerating the total number of eggs recovered. Positive *Ascaris* recovery controls were sampled immediately after application without the mopping procedure for the removal experiments. For survival experiments, positive *Ascaris* recovery controls were sampled immediately after application (time =0 days), and at each time point (2, 14, and 28 days) for the 34°C condition, 15°C condition, OPC mortar mix tiles, and OPC fly ash mortar mix tiles. Positive *Ascaris* recovery controls give insight into the total number of *Ascaris* eggs recovered from the tile surface after initial application given our washing method. The positive *Ascaris* recovery controls differ from the main study outcome because they measure the total number of eggs recovered, versus the number of viable or developed eggs. Negative controls were created by choosing a random quadrant on each cement mix tile prior to *Ascaris* inoculation. 2 mL of autoclaved DI water was added to one 9 cm^2^ quadrant of the tile and allowed to dry until the surface was no longer visibly wet (∼1 hour). The negative control was then recovered using the washing method and incubated in darkness for 32 - 35 days at 26°C. The positive and negative controls were enumerated as previously described.

The proportion of fertilized eggs in the stock was determined by spiking 25 μL of stock solution to 15 mL of autoclaved deionized (DI) water and 15 mL of 0.1 N sulfuric acid (Sigma-Aldrich, St. Louis, MO, USA). Samples were then immediately enumerated as previously described without incubation.

The number of control samples collected varies based on the sample type collected. The positive controls collected can be divided into three categories: stock controls without any incubation, application controls, and recovery controls. Five (5) measurements were taken for the stock controls without any incubation. Sixty-six (66) application controls were measured in total for the removal and survival experiments. Two-hundred and sixty (260) recovery controls were measured in total for the removal experiments; 52 for the removal conditions (26 trials x 2 cement mixes) and 208 for the survival conditions (13 trials x 4 time points x 2 temperature conditions x 2 cement mixes). Eighty (80) tiles were used throughout the experiments, and negative controls were measured for each tile.

### Data Analysis

Data analysis was performed with R (R: A Language for Statistical Computing, version 1.2.5042; R Foundation for Statistical Computing, Vienna, Austria) and G*Power [55].

The percent removal of *Ascaris* eggs was calculated using Equation 1:

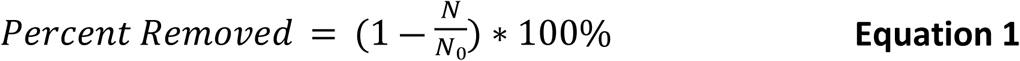

where *N* is the number of viable eggs after the mopping and *N_o_* is the initial number of viable eggs before mopping. The number of viable eggs for removal experiments was determined through the conventional method of viability assessment.

We assessed survival using percent viability and calculation of first-order decay rate constants. Calculations used both the number of viable eggs through the conventional method and the developmental evaluation method (referred to as “Developed” in the survival experiment results). We used Equation 2 to calculate percent viability:

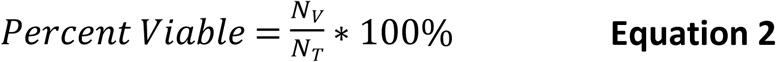

where *N_V_*is the number of viable at each time point and *N_T_* is the total number of eggs recovered at each time point.

To estimate the first-order decay rate constant (*k*), we fit the first-order log-linear regression with a normal family and identity link described in Equation 3:

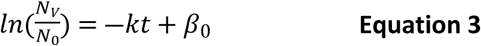

where *N_o_* is the initial number of viable eggs, *N_V_* is the number of viable eggs after a time of *t* (days) on the cement-based surface, *k* is the first-order decay rate constant (days*^−^*^1^), and *β*_0_ is the y-intercept. While some studies use log reductions with a lag [60,61], we did not observe a lag phase in our data (Figure S2). Models were fit for each experimental condition (4 regressions, 2 mix designs x 2 temperatures) as well as for each trial to obtain *k* for statistical analysis.

An unpaired, two-tailed t-test was used to determine differences in removal experiment results, while an ANOVA was used for differences in the *k* of the survival experiments. Interactions were not considered in our analysis. A Tukey honestly significant post-hoc test followed the ANOVA. Additional unpaired, two-tailed t-tests and ANOVAs with fixed effects were performed on subsets of data to determine differences in controls and variances over time. Data was tested for normality using Shapiro-Wilk tests and all analyses were performed using a significance level of *α* = 0.05.

### Literature Review of *Ascaris* Survival at Different Temperatures

In order to assess how our survival experiment results on cement-based surfaces compared to the survival of *Ascaris* eggs in other matrices, we conducted a systematized literature review of peer-reviewed articles broadly related to *Ascaris* survival in temperature conditions relevant to human habitats. A systematized literature review is similar to, but less stringent than, a systematic review [62]. The literature review included articles published up until December 16, 2024 using Web of Science, PubMed, Scopus, and Google Scholar. We used the search string (survival OR decay OR inactivation OR fate OR persistence OR viability) AND (“ascaris” OR roundworm) AND (temperature*) across titles, abstracts, and keywords in Web of Science, PubMed, and Scopus and incorporated all articles found. We used the search phrase “effect of temperature on ascaris” in Google Scholar and incorporated the first 50 articles generated. After duplicates were identified in Covidence (Covidence, Melbourne, Australia) and removed, 199 articles remained. Articles were then screened by one reviewer for the following requirements: (1) publication in English, (2) primary data collection (no reviews or modeling papers), (3) microscopy experiments with *Ascaris* eggs, (4) constant-temperatures experiments, (5) experimental (or control) temperatures less than 45°C, (6) experiments (or controls) without inactivation methods besides temperature, and (7) decay-rate constants, inactivation time, or reduction in viable *Ascaris* eggs reported.

We chose to include studies that investigated inactivation at temperatures less than 45°C because 45°C is at the upper bound of temperatures in which humans habitate; additionally, at temperatures below 45°C, *Ascaris* eggs are thought to have a different inactivation-temperature relationship than higher temperatures [60,63,64]. We also only included data from studies with low ammonia (no added ammonia and no studies in only urine) and where pH was typically neutral, as ammonia has been found to promote the inactivation of *Ascaris* and pH has had mixed effects [44,49,61,65–67]. Additionally, we only included results from aerobic studies, as floors in households are part of an aerobic environment and most studies agree that aerobic conditions accelerate inactivation [60,64].

After title/abstract screening of the initial 199 articles, 83 articles remained for full-text screening. After full-text screening, 26 papers were selected for data extraction. For five studies, data was approximated from figures using PlotDigitizer (PORBITAL). Time to 99% inactivation, when not reported by the study, was calculated assuming a log-linear inactivation model, even if a lag time was reported.

## Results

### Controls

We did not detect any *Ascaris* eggs in our negative controls (samples recovered from the cement-based tiles before *Ascaris* inoculation). For positive controls, the pre-incubation controls showed that over 98% of eggs were fertilized (single-celled) in the stock solution from 4°C storage. The application controls showed that 95.9% (SD = 1.9%) of eggs successfully larvated in the positive controls after incubation for 32 - 35 days at 26°C. Additionally, the mean number of total eggs applied to the cement-based tiles across experimental conditions was 510 eggs (standard deviation, SD = 106 eggs) across all trials and this was similar between mix designs (p = 0.87). The distribution of the total number of eggs applied to the tiles was normal, as confirmed by the Shapiro-Wilk test (p = 0.60) and visualized in a histogram (SI, Figure S3). In terms of recovery controls, the mean number of total eggs recovered from the tiles was 172 eggs (34% of applied eggs, SD = 65 eggs) and the number of total eggs recovered had a normal distribution (Shapiro-Wilk test p = 0.60).

### Experimental Setup

The dry weight of the solids in the stock volume controls yielded a mean dry weight of 3.26 mg for 0.25 µL of stock solution, yielding approximately 3.6 g/m² of solids in the experiments when spread over the tile quadrant.

### Removal Experiments

The mean removal of viable eggs by mopping for OPC fly ash mortar mix tiles was 95.2% (SD = 4.6%) and 95.9% (SD = 3.3%) for OPC mortar mix tiles (Figure 1). A t-test comparing OPC fly ash mortar mix tiles and OPC mortar mix tiles showed no significant difference in the removal efficiency of viable eggs by mopping between the groups (p = 0.51). Similarly, tile type did not significantly impact total *Ascaris* egg recovery for the removal experiments (p =0.10).

**Figure 1.**
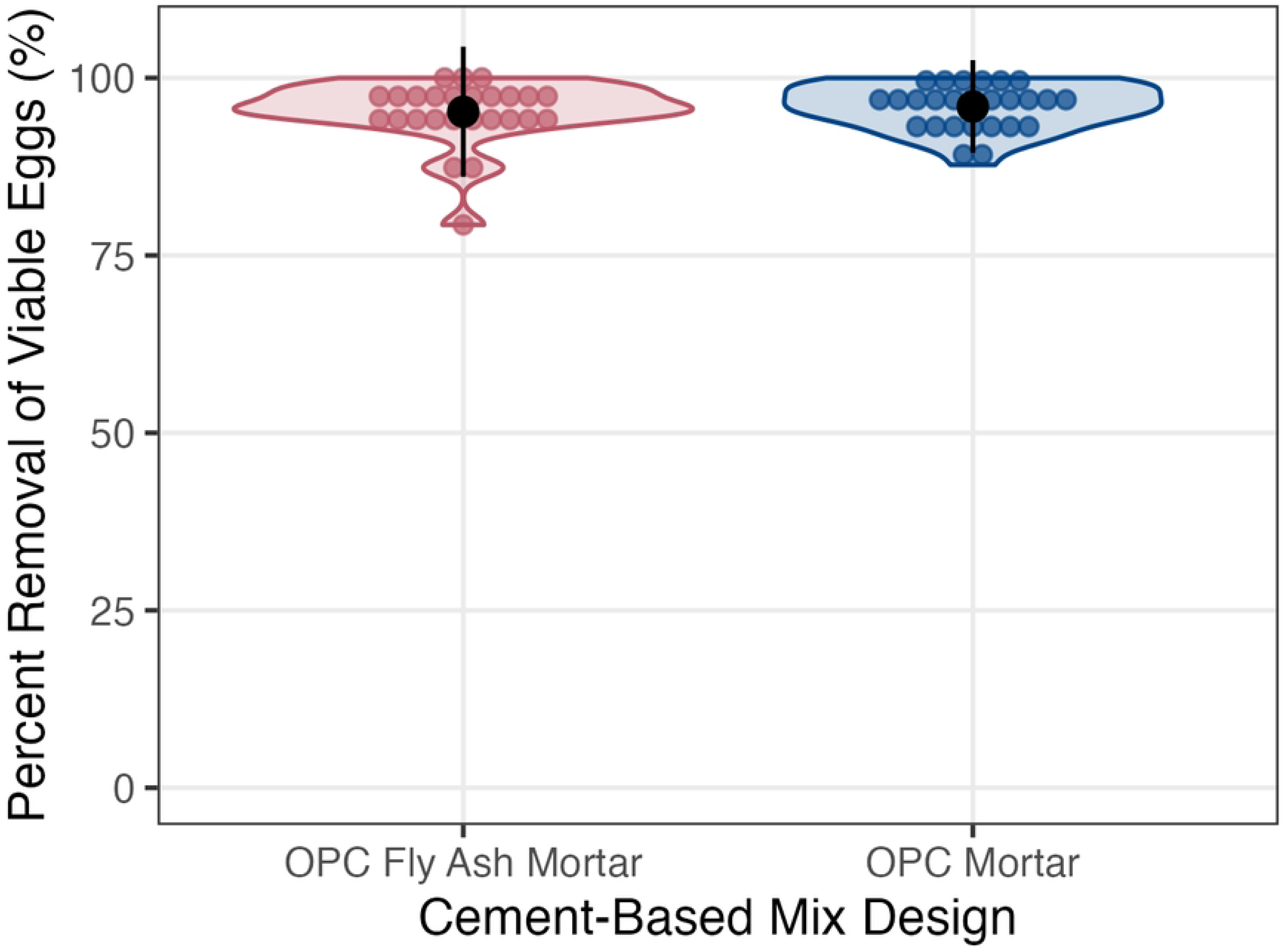

### Survival Experiments

The percent of viable eggs of total eggs (Equation 2) did not differ between the OPC fly ash mortar mix tiles and the OPC mortar mix tiles at the 0, 14, and 28-day time points (Figure 2, individual trial graphs available in the SI) for either enumeration method used. In contrast, at 2 days, percentages from both enumeration methods differed by mix design (p = 2.4 x 10^-3^ and p = 6.2 x 10^-4^, respectively). When the data were analyzed over the total length of the experiment rather than the individual time points, the *k* values from the linear regressions (Equation 3) of *Ascaris* egg survival for each trial (SI, Figure S2, S6, & S7) were similar between mix designs (p = 0.62). Across all survival experiments, the median *k* value was 0.029 day^-1^.

**Figure 2.**
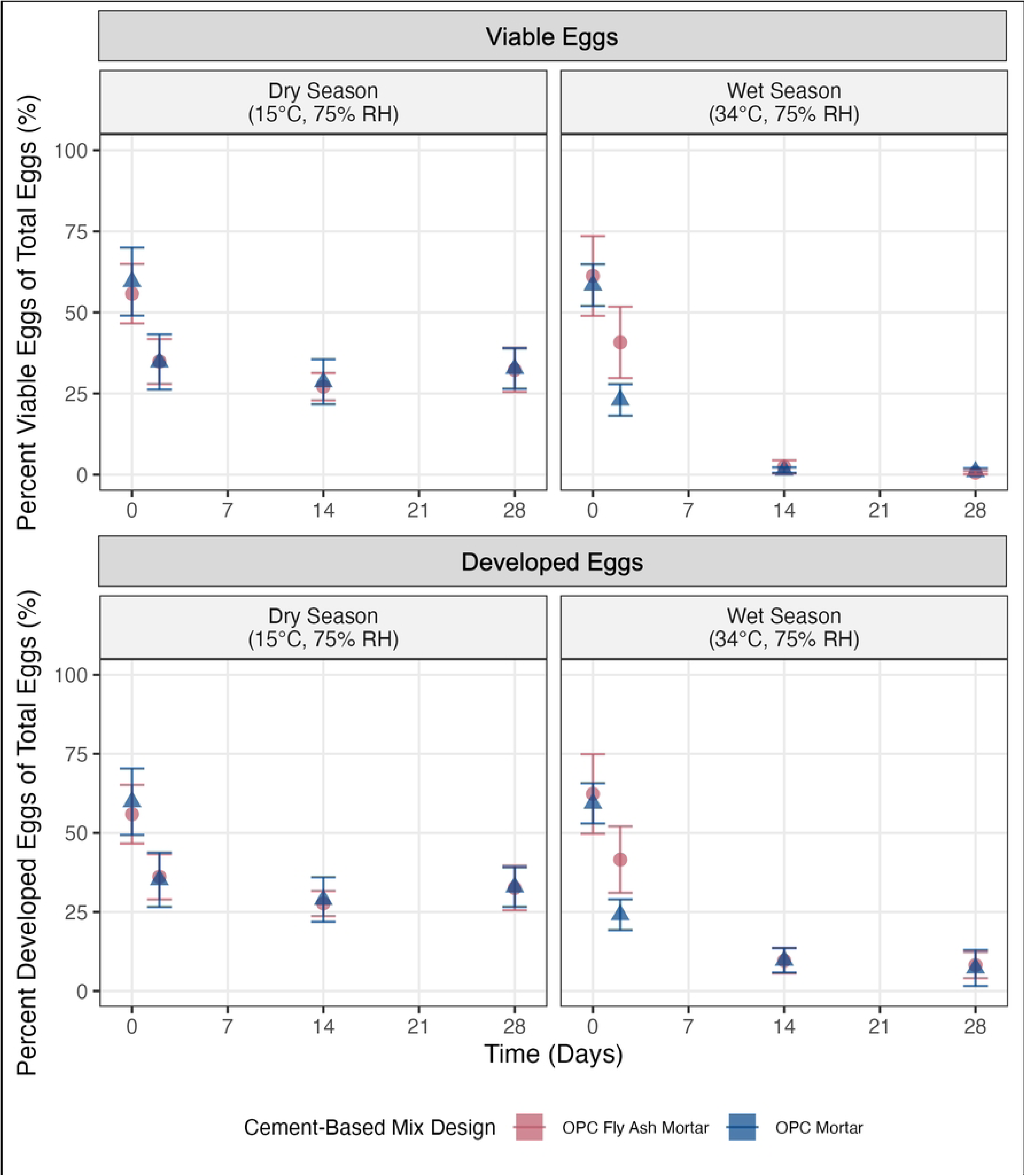

The *k* values (Table 1) varied significantly between temperatures (p = 4.2 x 10^-25^) and enumeration methods (p = 2.4 x 10^-8^). At 34°C, when all trials were combined, *k* values were 0.15 day^-1^ (p = 2.3 x 10^-20^) and 0.060 day^-1^ (p = 1.1 x 10^-11^) for conventional and developmental enumeration, respectively, on fly ash mortar tiles and 0.13 day^-1^ (p = 2.9 x 10^-17^) and 0.066 day^-1^ (p = 1.1 x 10^-12^) for conventional and developmental enumeration, respectively, on mortar tiles. These values indicate the inactivation of *Ascaris* eggs with time, as *k* values are significantly different from zero. In contrast, at 15°C, no significant differences in *k* from zero were observed. At 15°C, when all trials were combined, *k* values were −1.1 x 10^-3^ day^-1^ (p = 0.82) and −7.9 x 10^-4^ day^-1^ (p = 0.87) for conventional and developmental enumeration, respectively, on fly ash mortar tiles and 1.4 x 10^-3^ day^-1^ (p = 0.80) and 1.5 x 10^-3^ day^-1^ (p = 0.78) for conventional and developmental enumeration, respectively, on mortar tiles. After 28 days, there was a mean reduction of eggs quantified using the conventional enumeration method of 59.2% at 34°C and 25.5% at 15°C, across trials and cement-based mix designs. Post-hoc tests confirm that wet season, higher temperature *k* values exceed dry season, low-temperature *k* values (p = 1.2 x 10^-10^, mean difference of 0.11 day^-1^ in *k* between conditions). Post-hoc tests also confirm that *k* values obtained using the conventional method counts had a greater magnitude (indicating faster inactivation in the same time period) than those obtained from developed egg counts (p = 2.3 x 10^-8^, mean difference of 0.05 day^-1^ in *k* between enumeration methods).

**Table 1:**
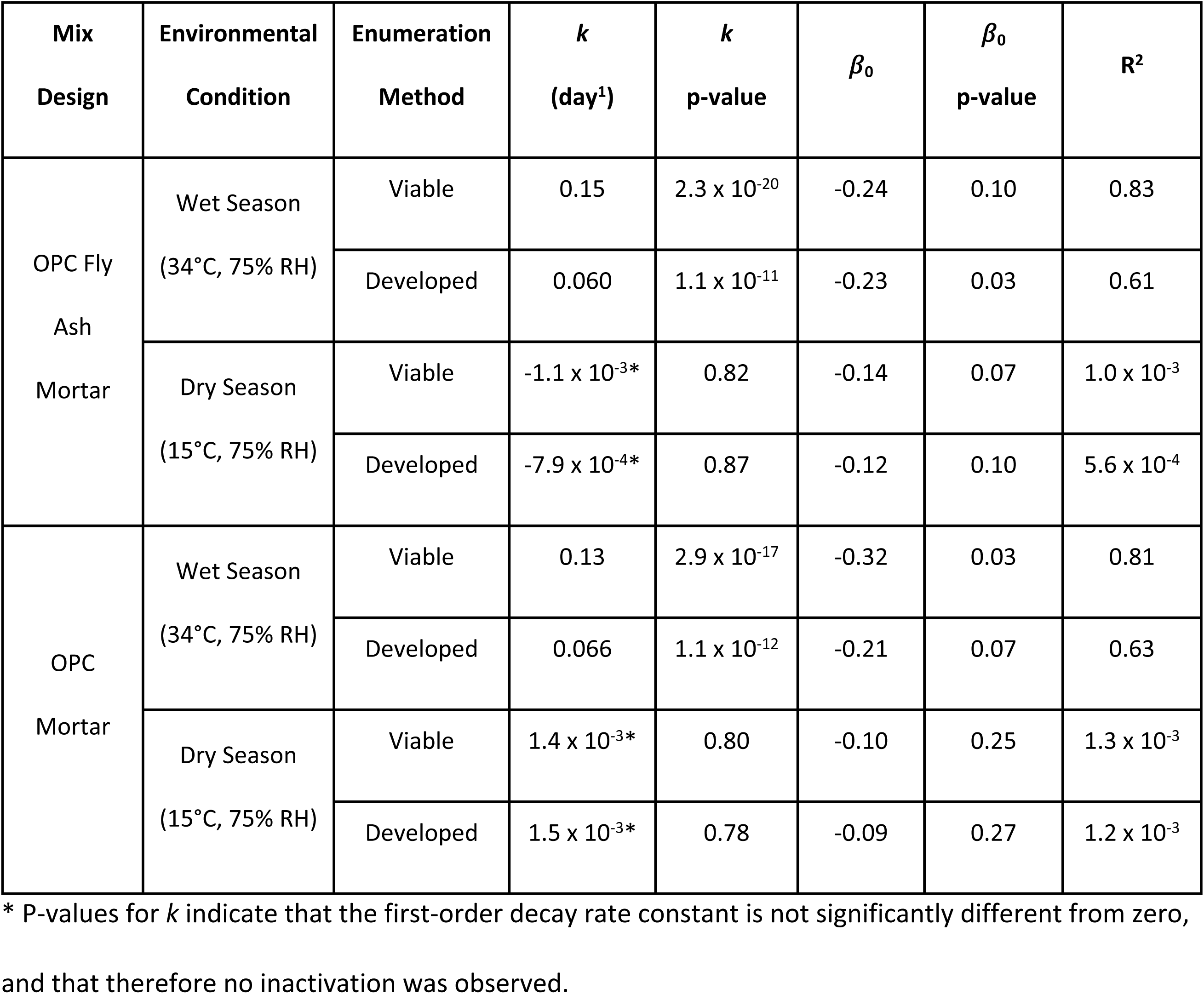
Regression results, including first-order decay rate constants (*k*) and y-intercept values (*β*_0_) for each cement-based mix design, experimental condition, and enumeration method used.

| Mix Design | Environmental Condition | Enumeration Method | $k$ (day<sup>-1</sup>) | $k$ p-value | $\beta_0$ | $\beta_0$ p-value | R<sup>2</sup> |
| --- | --- | --- | --- | --- | --- | --- | --- |
| OPC Fly Ash Mortar | Wet Season<br>(34°C, 75% RH) | Viable | 0.15 | $2.3 \times 10^{-20}$ | -0.24 | 0.10 | 0.83 |
| | | Developed | 0.060 | $1.1 \times 10^{-11}$ | -0.23 | 0.03 | 0.61 |
| | Dry Season<br>(15°C, 75% RH) | Viable | $-1.1 \times 10^{-3*}$ | 0.82 | -0.14 | 0.07 | $1.0 \times 10^{-3}$ |
| | | Developed | $-7.9 \times 10^{-4*}$ | 0.87 | -0.12 | 0.10 | $5.6 \times 10^{-4}$ |
| OPC Mortar | Wet Season<br>(34°C, 75% RH) | Viable | 0.13 | $2.9 \times 10^{-17}$ | -0.32 | 0.03 | 0.81 |
| | | Developed | 0.066 | $1.1 \times 10^{-12}$ | -0.21 | 0.07 | 0.63 |
| | Dry Season<br>(15°C, 75% RH) | Viable | $1.4 \times 10^{-3*}$ | 0.80 | -0.10 | 0.25 | $1.3 \times 10^{-3}$ |
| | | Developed | $1.5 \times 10^{-3*}$ | 0.78 | -0.09 | 0.27 | $1.2 \times 10^{-3}$ |

\* P-values for $k$ indicate that the first-order decay rate constant is not significantly different from zero, and that therefore no inactivation was observed.

The surface roughness of the two different mix designs could influence the recovery of the total number of *Ascaris* eggs. Similarly, *Ascaris* eggs could desiccate and lose adherence properties with time on cement-based surfaces or with high temperatures, which would also affect recovery. To explore these factors, we conducted an ANOVA to test the effects of tile type, time, and environmental conditions on recovery. The average recovery of *Ascaris* eggs from the tiles across all conditions was 34% of the total applied eggs. Tile type did not significantly impact the recovery of total *Ascaris* eggs from the tiles (p = 0.17), but both time (p = 5.7 × 10⁻⁹) and temperature (p = 1.2 × 10⁻⁵) significantly influenced recovery in survival experiments. Overall, recovery increased from an average of 128 eggs at time 0 (25% of the total applied eggs) to 202 eggs at 28 days (40% of the total applied eggs). At the low temperature, an average of 156 eggs were recovered (31% of the total applied eggs), whereas in the high temperature, the mean recovery was 194 eggs (38% of the total applied eggs) (Figure S4). We have included all recovery percentages, as well as a correction for *k* for recovery differences over time, in the SI. The correction resulted in slightly increased k values (mean difference of 0.014 day^-1^; median *k* value of 0.042 day^-1^ with the correction versus 0.029 day^-1^ using unadjusted data). In the main text, we report unadjusted decay-rate constants as a conservative estimate of *Ascaris* egg inactivation.

### Literature Review of *Ascaris* Survival at Different Temperatures

Across 27 studies (including this study), the most common matrix for measuring *Ascaris* egg survival was fecal sludge and only this study provided survival estimates on solid surfaces [43,44,60,61,64,68–88]. Matrices and study types were simplified for visualization in Figure 3. The complete data set and visualization are available in the SI. Temperatures investigated ranged from 0 to 44°C, and 105 unique data points of time to 99% inactivation were reported or derived from literature data. Most (23/26) studies from the literature used the conventional method of enumeration. To compare our results to these studies, we used the results of our study obtained with the conventional method of enumeration. In our study, to achieve 99% inactivation of *Ascaris* eggs on cement-based surfaces, an inactivation time of 29 to 33 days is needed for surfaces at an ambient temperature of 34°C. In contrast, over 3000 days (8.8 years) is required to achieve 99% inactivation of *Ascaris* eggs on cement-based surfaces at 15°C.

**Figure 3.**
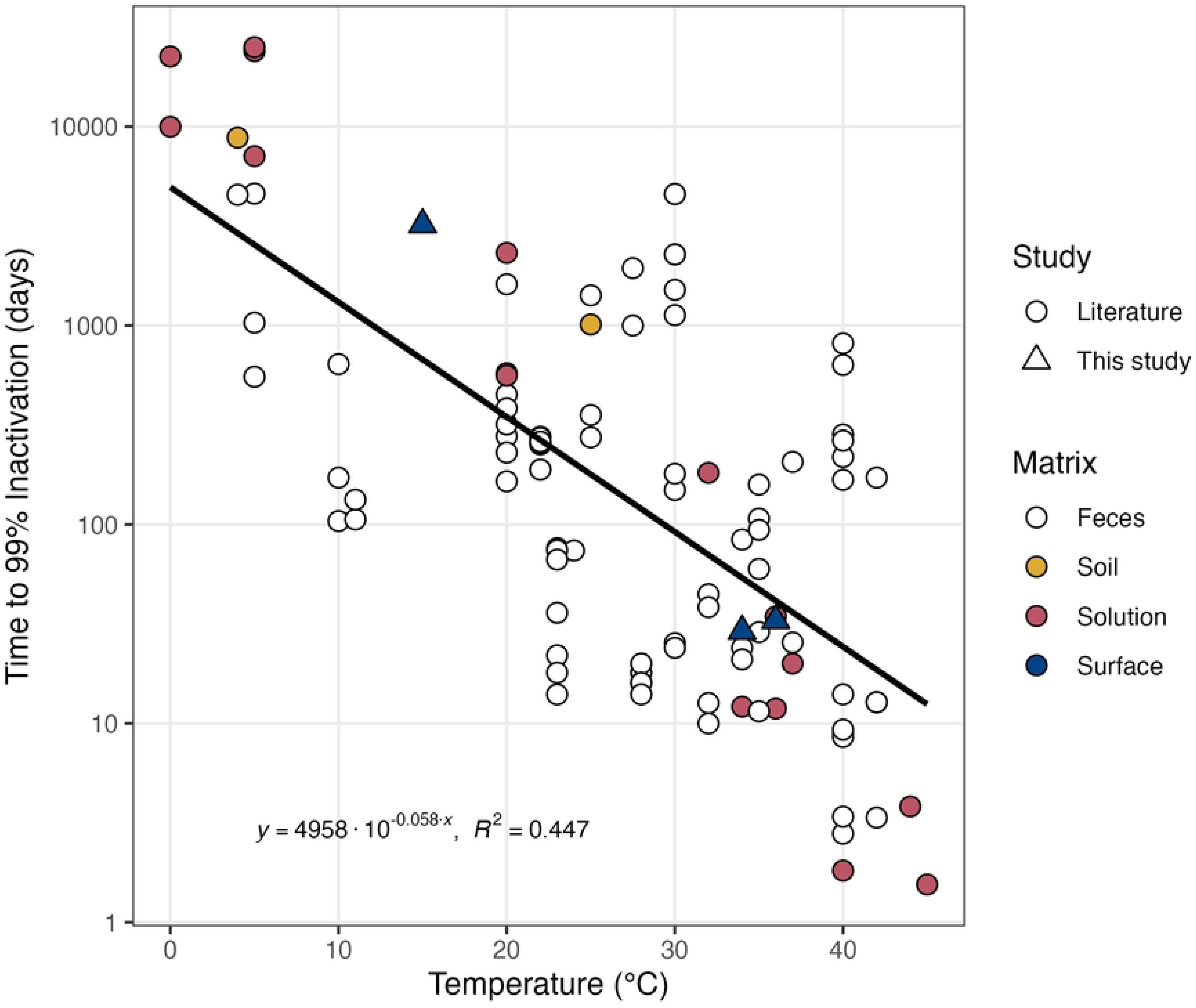

Generally throughout the studies, at higher temperatures, the time to 99% inactivation of *Ascaris* eggs was lower. This relationship was observed across all matrices. From the data set of the literature and the data obtained in this study, we approximated the time-temperature relationship (Equation 4) for *Ascaris* egg inactivation at temperatures relevant to human environmental exposures to be:

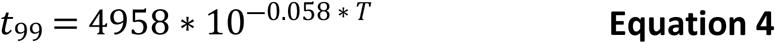

where *t_99_* is the time for 99% inactivation in days and *T* is the temperature in °C. This regression has an R^2^ value of 0.45. Using Equation 3, we estimate a general time of 99% inactivation of *Ascaris* eggs of 669 days (1.8 years) at 15°C and 53 days at 34°C.

## Discussion

We found that Ascaris survival and removal from cement surfaces were similar between different cement mixes, including a mix with lower embodied carbon. These findings support the use of these mixes in the ongoing randomized control trials to measure health impacts. The study outcomes indicate that mopping is a viable cleaning method for Ascaris and that the time to inactivation is longer at cooler temperatures, suggesting that cooler temperatures may be an important condition in which to focus intervention efforts.

The mean removal percentage of *Ascaris* eggs from mopping was above 92% for both cement-based mix designs tested. Overall, the removal percentage was lower than reported in prior studies, which investigated removal in liquid matrices, with removal methods including adding ammonia, using 254-nm ultraviolet light, and filtering [42–44]. While these experiments often observed high inactivation or removal rates of *Ascaris* eggs (+99% under some conditions), the results of these studies are not directly applicable to the removal of *Ascaris* from surfaces, especially in LMICs, where many chemical products are not attainable for rural households. Thus, while the removal efficiency from mopping (+92%) is lower than that achieved with chemical disinfectants, it is more applicable to low-income, rural settings where access to chemical products is limited. A limitation of this study is that while mopping removed a significant proportion of *Ascaris* eggs from cement-based tile surfaces, the eggs may not have been inactivated. Viable eggs may persist on cleaning tools, such as mops, which could lead to cross-contamination. This phenomenon was observed in one previous study which investigated the disinfection of glass and plastic surfaces and found that while eggs were removed from surfaces, they were not inactivated on the cleaning tool [89]. Although additional passes with a cleaning tool could potentially enhance removal efficiency, additional passes may also increase the likelihood of eggs adhering to the cleaning tool and being redistributed elsewhere within the household. Future studies could investigate *Ascaris* egg survival on cleaning tools, redistribution of eggs elsewhere in the home, and potential disinfection methods for cleaning tools.

In terms of *Ascaris* egg survival on cement-based surfaces, our results showed that *k* did not differ by cement mix design, but were influenced by temperature. Specifically, at 34°C, *k* had a greater magnitude, indicating faster inactivation, than at 15°C. Moreover, *k* at 15°C was not significantly different from zero, indicating no inactivation was observed during the dry season conditions. A low *k* at low temperatures for human habitats agrees with previous studies, which found that the *k* values of *Ascaris* eggs largely depended on temperature and oxygen availability. In previous studies, the *k* values increased in magnitude with temperature at a predictable rate, and at temperatures above 60°C, eggs were inactivated within a few minutes [63,90].

Our findings are consistent with *Ascaris* survival trends in different environmental matrices reported in other studies. When comparing our results to studies selected from our literature review with inactivation rates in realistic environmental conditions (<45°C), results suggest that the time to 99% inactivation of *Ascaris* eggs on cement-based surfaces is similar to liquid and semi-solid matrices, especially at 34°C (Figure 3 and SI). Similar inactivation times of *Ascaris* eggs between cement-based surfaces and liquid and semi-solid matrices indicate that replacing soil floors with cement-based flooring may not inherently reduce *Ascaris* egg concentrations, as there is no increased inactivation on the cement-based surface. Instead, decreased *Ascaris* egg concentrations on cement-based surfaces may be a result of the ease of cleaning the surface, as discussed in the mopping experiments.

The *k* values reported for *Ascaris* eggs in this study may be impacted by the high concentration of total solids present in the spike solution (3.6 g/m²), which exceeds typical field levels reported in rural Bangladeshi households with cement-based floors (0.2 g/m², SD=0.3, of dust weight from sweeping) [91]. The solids likely consist mostly of organic matter, and organic matter like fecal matter has been shown to protect *Ascaris* eggs from environmental stressors [60], suggesting that our experimental conditions may have overestimated egg survival compared to typical household scenarios. Additionally, some previous studies have observed a lag phase of up to 12 weeks [60,61] in the inactivation of *Ascaris,* which we may have observed under the conditions described herein if we sampled for a longer time.

The choice of enumeration method significantly affected inactivation estimates; conventional enumeration yielded larger *k* values compared to the developmental enumeration method. This discrepancy indicates that conventional methods may underestimate viable egg numbers by excluding late developmental stages and is in line with hypotheses from previous studies [46,52,54]. The resultant difference depended on the temperature of the survival experiments. In higher temperature conditions, the difference between the *k* values calculated from conventional and developmental enumeration methods was greater. Additionally, at the higher temperature, the linear decay model has increased R^2^ values, indicating it fits better for the conventional enumeration versus the developmental enumeration method. These results emphasize that while the conventional method of enumeration simplifies decay modeling, it may provide an incomplete picture of *Ascaris* egg survival.

The recovery percentage of *Ascaris* eggs from cement-based surfaces in this study was approximately 34% and did not vary by cement mix, although variations were observed across all samples under different temperatures and times. Changes in recovery based on temperature and time are likely due to changes in egg morphology, such as desiccation or alterations in surface adhesion properties. Recovery differences over time could have implications for survival and removal scenarios in the field. For instance, eggs from fresh fecal contamination on cement-based flooring may be less easily removed from cement-based surfaces. With time, dried eggs may become less adherent to the cement-based surfaces, facilitating their removal by mopping but also increasing their potential for dispersal and transmission.

Overall, this study contributes novel insights into both *Ascaris* egg removal and survival dynamics on surfaces, providing some of the first estimates of removal and *k*. Repeated trials (26 per mix design for removal experiments and 13 per condition for survival experiments) strengthened our results. Results indicate that cement-based flooring may reduce *Ascaris* egg concentrations due to easier cleaning methods, but survival was similar to other matrices. By conducting experiments with both a traditional and a more sustainable cement-based flooring option, our results show that *Ascaris* eggs behaved similarly on both surfaces in terms of removal and survival. Our results highlight that the sustainable OPC fly ash mortar mixes can serve as an effective alternative to traditional cement-based flooring in disease interventions, as both have potential strengths to reduce *Ascaris* transmission in rural LMIC households.

## Data Availability

Anderson, C. E., Hanif, S., Hernandez, J., Crider, Y., Lepech, M., Billington, S. L., Boehm, A. B., and Benjamin-Chung, J. (2025). Ascaris suum Survival and Removal Code and Complete Data Set. Version 2. Stanford Digital Repository. Available at https://purl.stanford.edu/rj162zg5361/version/3. https://doi.org/10.25740/rj162zg5361.

## Acknowledgments

This work was supported by a grant from the Stanford Woods Institute for the Environment (281131) to JBC, the NSF Graduate Research Fellowships Program (GRFP) to CEA, and Air Force Institute of Technology Fellowship Program and John A Blume Earthquake Engineering Center Fellowship at Stanford University to JH. JBC is a Chan Zuckerberg Biohub Investigator. This study was performed on the ancestral and unceded lands of the Muwekma Ohlone people. We pay our respects to them and their Elders, past and present, and are grateful for the opportunity to live and work there.

